# Dissecting the biological impact of *GBA1* mutations using multi-omics in an isogenic setting

**DOI:** 10.1101/2025.06.12.656971

**Authors:** Pilar Álvarez Jerez, Peter A. Wild Crea, Dhairya Patel, Logan M. Glasstetter, Christiane Alvarez, Mary B. Makarious, Erika Lara, Yu Chen, Caroline B. Pantazis, Kimberly Paquette, Laksh Malik, Mike A. Nalls, Xylena Reed, Andrew B. Singleton, Kimberley J. Billingsley, Justin A. Mcdonough, Gang Ning, William C. Skarnes, Mina Ryten, Ellen Sidransky, Mark R Cookson, Sasha Beilina, Cornelis Blauwendraat

## Abstract

*GBA1* is a risk gene for multiple neurodegenerative diseases, including Lewy Body Dementia and Parkinson’s disease, and biallelic pathogenic variants in the gene result in the lysosomal storage disorder Gaucher disease. *GBA1* encodes the enzyme glucocerebrosidase (GCase), and alterations in the gene result in reduced enzymatic activity, which affects lysosome function downstream. Induced pluripotent stem cells (iPSCs) are a useful tool for testing the functional consequences of gene variants in an isogenic setting. Additionally, they can be used to perform multiomic studies to explore biological effects independent of disease mechanisms. Using CRISPR-edited isogenic KOLF2.1J iPSC lines containing pathogenic *GBA1* variants D409H (p.D448H), D409V (p.D448V) and GBA1 knockout line generated by the iPSC Neurodegenerative Disease Initiative (iNDI), we examined potential molecular mechanisms and downstream consequences of GCase reduction. In this study, we confirm that this isogenic series behaves as expected for loss of function variants, despite the known difficulties with *GBA1* editing. We identified that there are limited overlapping results across cell types suggesting potential different downstream effects caused by *GBA1* variants. Additionally, we note that RNA-based quantitation may not be the best method to characterize GCase mechanisms, but protein and metabolomic analyses may be used to evaluate differences across genotypes.

## Introduction

Gaucher disease (GD) is an autosomal recessive lysosomal storage disorder caused by biallelic pathogenic variants in *GBA1*, which encodes the enzyme glucocerebrosidase (GCase) ^1^. GCase is essential for the breakdown of glucocerebroside, a lipid component in cell membranes. Loss of function variants in *GBA1* lead to insufficient enzyme activity, resulting in the accumulation of glucocerebroside within lysosomes ^2^. The disease is clinically divided into three main types: non-neuronopathic (type 1), acute neuronopathic (type 2), and chronic neuronopathic (type 3) ^3^.

Additionally, *GBA1* variants have been associated with an increased risk of Parkinson’s disease (PD) and Lewy Body Dementia (LBD), which are complex neurodegenerative diseases presenting with tremors, dystonia, bradykinesia, and dementia ^4,56^. Over 400 different mutations in *GBA1* have been identified that lead to GD, and some are also linked to increased PD risk when in a heterozygous state, with odds ratios varying from 1.5 to ∼7 ^7^. Some of the more common pathogenic *GBA1* variants include p.N370S (p.N409S), which leads to type 1 GD, increased PD risk, and has increased prevalence in the Ashkenazi Jewish population, and p.L444P (p.L483P), which is associated with increased PD risk and when homozygous generally leads to type 3 GD ^8^. Other common variants, such as p.E326K (p.E365K) and rs3115534-G confer PD risk but do not cause GD ^9,10^.

The use of induced pluripotent stem cell (iPSC) isogenic lines can offer numerous advantages for studying disease biology related to specific variants compared to population-level studies. By eliminating genetic background variability, isogenic lines ensure that phenotypic differences can be attributed solely to the variant of interest, allowing, for example, different alleles to be directly compared to each other in the context of a shared genetic background ^11^. Here, we use an isogenic model harboring two GD-causing mutations, p.D409H (p.D448H, rs1064651) and p.D409V (p.D448V, rs77369218) ^12,13^, and a *GBA1* knockout to explore downstream effects of these variants on a multiomics level. p.D409H is a rare *GBA1* variant that has been reported with an allele frequency of ∼0.00012 in the population (https://gnomad.broadinstitute.org/gene/ENSG00000177628?dataset=gnomad_r4), while p.D409V is a very rare mutation reported in a heterozygous state in several GD patients and commonly used in mouse models ^14^.

Here, we use CRISPR-edited isogenic lines to look for potential molecular mechanisms of *GBA1* and additional affected genes in GCase loss of function-based disease. The main motivation is that any gene, protein, or metabolite correlated with GCase activity could be a potential interactor and provide valuable information regarding the mechanism underlying *GBA1* associated PD. We differentiated these isogenic cell lines into forebrain cortical neurons and microglia, two potentially disease-relevant brain cell types, and performed RNAseq, proteomic, and metabolomic evaluations to assess differences across genotypes. Using this data we explored the question: are effects of *GBA1* variants similar or dissimilar in these two important brain cell types? Our results validate the use of this isogenic series and suggest that protein and metabolomic read-outs are potentially more useful to measure downstream effects of *GBA1* variants compared to RNA based methods. In addition, we lay the foundation for potential follow-up larger scale studies exploring the downstream effects of *GBA1* mutations.

## Methods

### Induced pluripotent stem cell line selection

For this study, we used the KOLF2.1J cell line, which is employed by the iPSC Neurodegenerative Disease Initiative (iNDI) at the NIH’s Center for Alzheimer’s and Related Dementias (CARD) as a high-quality reference cell line to use in neurodegenerative research ^11^. This line was selected after deep characterization of genomic status, functional characteristics, and differentiation potential of multiple iPSC lines, of which KOLF2.1J performed the best. The KOLF2.1J parental line and derivatives can be found here https://www.jax.org/jax-mice-and-services/ipsc. CRISPR-Cas9 editing on the KOLF2.1J wild type IPSC line was performed as described in Skarnes et al. 2019 ^15,16^ to generate variants of interest (p.D409H and p.D409V). p.D409H is a known cause of GD in the homozygous state and increases PD risk in its heterozygous state. p.D409V appears to be likely pathogenic although only heterozygous cases have been reported. In addition, a homozygous *GBA1* knockout cell line was generated by removing 5.4 kb containing exons 1-9 with Cas9 RNP with the addition of a 100-mer bridging oligo spanning the deletion junction. Guide and primer sequences can be found in Supplementary Table 1.

For all cell culture experiments, we included three replicates of WT and three separate clones from each CRISPR edited genotype line, leading to a total of 18 iPSC lines (WT x3, D409H_heterozygous x3, D409H_homozygous x3, D409V_heterozygous x3, D409V_homozygous x3, knockout x3).

#### Induced pluripotent stem cell line differentiation

To investigate the effects of *GBA1* mutations in additional cell types, the KOLF2.1J parental and KOLF2.1J-GBA1 edited iPSC lines were differentiated into forebrain cortical neurons and microglia. Cells were differentiated into forebrain cortical neurons following the protocol described in Blauwendraat et al 2020 ^17^. In short, iPSCs were grown in E8 media (ThermoFisher, A1517001) on Matrigel until 90% confluent after which they were transitioned to N3 media (50% DMEM/F12, 50% Neurobasal™ with 1× penicillin-streptomycin, 0.5× B-27™ minus vitamin A, 0.5× N2 supplement, 1× GlutaMAX™, 1× NEAA, 0.055 mM 2-mercaptoethanol and 1 μg/ml Insulin) plus 1.5 μM dorsomorphin (Tocris Bioscience, 3093) and 10 μM SB431542 (Stemgent, 04-0010-base) as previously reported ^18^. Media was replaced every day for 11 days and cells were fed with additional N3 media. On day 20, the differentiating cells were split and were then fed daily with N4 media (same as N3 plus 0.05 μM retinoic acid, 2 ng/ml BDNF and 2 ng/ml GDNF) until day 26 after which they were frozen. Differentiation was confirmed by immunocytochemistry for neuronal markers β-III tubulin (Novus Biologicals, NB100-1612) and MAP2 (Santa Cruz Biotechnology, sc-20172) with nuclei counterstained for Hoechst 33342 (ThermoFisher Scientific, H3570).

Microglia were differentiated as reported in Brownjohn et al 2018 ^19^. This protocol followed an established method for the derivation of primitive macrophage precursors (PMPs) as a starting point for microglia differentiation ^20,21^. In brief, iPSCs were passaged to single cells with TrypLE Express (Gibco, 12604013) and plated in 100uL embryoid body medium (EBM) (10 μM ROCK inhibitor, 50 ng/mL BMP-4, 20 ng/mL SCF, and 50 ng/mL VEGF-121 in E8). The following day, 80 µL of medium was removed from each well and replaced with 100 µL of EBM without ROCK inhibitor. EBs were cultured for an additional 2 days, with daily media changes using fresh EBM without ROCK inhibitor. Then, ∼200 embryoid bodies were transferred into a T75 flask and cultured in 20 mL hematopoietic medium (2 mM GlutaMax, 100 U/mL penicillin, 100 μg/mL streptomycin, 55 μM β-mercaptoethanol, 100 ng/mL M-CSF, and 25 ng/mL IL-3 in X-VIVO 15 [Lonza, LZBE02-060F]), after which medium was exchanged every 2 days. Microglia progenitor cells (MPCs) typically appeared between Day 12 and Day 18. The cells were collected by centrifugation, and approximately 4 million cells per flask were seeded into non-coated 75 cm² flasks for final differentiation in Microglia Maturation Media (MMM), consisting of Advanced RPMI, 2 mM GlutaMax™, 100 ng/mL IL-34, and 10 ng/mL GM-CSF. Cells were further differentiated for 10 additional days, with full MMM media changes every other day. After differentiations, we had three clones per genotype as with the parental iPSCs for a total of 18 lines of forebrain neurons and 18 lines of microglia, for a total of 54 samples across the three cell types.

### RNA extraction

RNA was extracted from all cell lines (5e6 cells per pellet) using the RNA Direct-zol miniprep kit and its corresponding protocol (Zymo Research, R2050, https://files.zymoresearch.com/protocols/_r2050_r2051_r2052_r2053_direct-zol_rna_miniprep.pdf). In short, samples were resuspended in 600 uL of TRI reagent and an equal volume of 100% ethanol. Each mixture was then transferred into a Zymo-Spin IICR Column, centrifuged, and transferred to a new collection tube. DNase I treatment was performed using the recommended guidelines. Zymo Research’s Direct-zol RNA PreWash and RNA Wash Buffer were used for wash steps. Finally, RNA was eluted in RNase-free water and was quality controlled on Agilent’s TapeStation 4200. All cell lines had a RIN > 8 (see **Supplementary Table 2** for additional details).

### GCase activity measurements assay

GCase enzymatic activity was determined based on the cleavage of the fluorogenic substrate 4-methylumbelliferyl-β-D-glucopyranoside (4-MUG), as previously described ^22,23^. In brief, freshly-prepared GCase buffer [McIlvaine buffer (0.2 M Na2HPO4 and 0.1 M citric acid titrated to pH 5.4); 0.25% (v/v) Triton X-100 (Cat.#T9284; Sigma-Aldrich); Roche cOmplete™, Mini, EDTA-free Protease Inhibitor Cocktail (Cat.#11836170001; 1 tablet/10 mL)] was activated with 0.2% (w/v) sodium taurocholate (Cat.#86339; Sigma-Aldrich). Protein was extracted from iPSC and neuron cellpellets (4x6 cells per pellet) in GCase buffer through pipetting, at least two 10-sec pulsed bath sonication steps (S-4000, Misonix), and 3 freeze-thaw cycles. The lysates were centrifuged for 15 min (21,000 x *g*; 4°C) and the supernatant containing extracted protein was collected. Protein concentration was measured via the Pierce™ BCA Protein Assay Kit (Thermo Scientific) and samples were diluted to 1 µg/µL with GCase buffer^22^

The 4-MUG GCase activity assay was conducted in a Greiner black 384-well plate (Cat.#781209), to which protein samples were added (10 µL/well). To correct for background noise, GCase buffer (5 µL/well) containing either conduritol B epoxide (CBE; 0.8 mM) or vehicle (DMSO, 8% v/v) was added. The plate was sealed, centrifuged, and incubated in a shaking plate incubator at 37°C and 600 rpm for 15 min. After brief centrifugation, 15 µL of assay buffer (2.5 mM 4-MUG; 0.25% v/v DMF in GCase buffer) was added to each well. The plate was sealed, centrifuged, and incubated at 37°C for 60 min while shaking at 450 rpm. After incubation, the plate was centrifuged and 30 µL of 1 M glycine stop solution (pH 10.5) was added to each well. Fluorescence was top read on a FlexStation® 3 Multi-Mode Microplate Reader (Molecular Devices). For each sample, GCase activity was calculated by subtracting the average fluorescence readout of CBE-inhibited wells from that of DMSO-control wells.

GCase differences across groups were calculated using a two-sided t-test in R for every genotype group against the wild type. Significance was determined by p < 0.05.

### RNA sequencing data generation

Illumina short-read RNA sequencing data was generated for all lines. RNA was extracted as described above, and library preparation and sequencing was completed by Psomagen Inc (https://www.psomagen.com/). Per Psomagen, the rRNA was removed, and the remaining total was fragmented and primed for cDNA synthesis via TruSeq Stranded Total RNA Library Prep kit reagents (96 Samples, Illumina #20020597), and incubated for 8 minutes at 95°C (C1000 Touch Thermal Cycler). The cleaved and primed RNA was reverse transcribed into first strand cDNA using SuperScript II Reverse Transcriptase (#18064-014, Thermo Fisher Scientific, Waltham, MA, USA); Actinomycin D and FSA (First Strand Synthesis Act D Mix) were added to enhance strand specificity. The second strand was synthesized using the 2nd strand master mix from the same TruSeq Stranded Total RNA kit (16°C incubation for 1 hour). To enable adapter ligation, the double-stranded cDNA (dscDNA) was adenylated at the 3’ end and RNA adapters were subsequently ligated to the dA-tailed dscDNA. Additional amplification steps were carried out to enrich the library material. The final library size was measured (D1000 ScreenTape, Agilent Tapestation System) and quantified (Quant-iT PicoGreen dsDNA Assay Kit). The sequencing library was then loaded onto a flow cell. Following cluster generation, the Illumina Sequencing by Synthesis (SBS) technology (Illumina Novaseq 1.5 5000/6000 S4 Reagent Kit-300 cycles [#20028312]; Illumina Novaseq 1.5 Xp 4-Lane Kit [#20043131]) was used to accurately sequence each base pair. Real Time Analysis software (RTA v3) was used to base-call data from raw images generated by the Illumina SBS technology. Binary BCL/cBCL files were then converted to FASTQ files using bcl2fastq (bcl2fastq v2.20.0.422)–an Illumina provided package.

### Metabolomics data generation

To assess the metabolomics profile of our samples, pellets (∼4-5e6 cells per pellet) were sent to Metabolon for metabolomics data generation. Per Metabolon Inc., the samples were homogenized and subjected to methanol extraction, then split into aliquots. Analysis of these aliquots was performed by ultrahigh performance liquid chromatography/mass spectrometry (UHPLC/MS) in the positive (two methods) and negative (two methods) mode. Metabolites were then identified by automated comparison of ion features to a reference library of chemical standards followed by visual inspection for quality control,as previously described in Dehaven et al. 2010. ^24^). Any missing values were assumed to be below the limits of detection andwere imputed with the compound minimum (minimum value imputation). Statistical tests were performed in ArrayStudio (Omicsoft) to compare data between experimental groups, with a significant threshold of p < 0.05. An estimate of the false discovery rate (Q-value) was also calculated to take into account the multiple comparisons that normally occur in metabolomic-based studies, with q < 0.05 used as an indication of high confidence in a result. Results were also normalized by protein values as measured by Bradford assay.

### Somalogic data generation

Protein lysates for each cell type were sent to the NIH Center for Human Immunology, Inflammation, and Autoimmunity to be run on a SomaScan assay. Per Somalogic, profiling of the proteome was carried out using the SomaScan® Assay, which is a highly multiplexed aptamer-based proteomic technology capable of making over 7,000 protein measurements. These chemically modified aptamers, called SOMAmer® Reagents, form complex three-dimensional shapes which bind to epitopes on their target protein with high affinity and specificity. The amount of the available protein epitope is read out by hybridizing the SOMAmer reagents in the SomaScan Assay eluate to complementary sequences on a DNA microarray. The fluorescence is measured as relative fluorescence units (RFU). The RFU readout is normalized and delivered in tab-delimited text files with the .adat extension.

### RNA sequencing data processing and analysis

Illumina FASTQ data were aligned to hg38 using STAR (v2.7.10) and salmon (v 1.10.1) was used to quantify gene expression. After salmon quantification, we generated a gencode gene map from the gencode v43 transcripts FASTA (https://www.gencodegenes.org/human/release_43.html) to generate a clean transcript name to gene name table. Lastly, we used the tximport ^25^ package in R to create a matrix with the salmon gene quantifications for each sample. Genes with an average transcripts per million (TPM) < 2 across all samples were removed from the analysis. This same process was done to generate matrices for all three cell types. Principal components (PC) were calculated using prcomp and plotted with autoplot in R for each cell type. We removed any gene where there was no variance across all entries, as those interfered with the principal component analysis (PCA). No additional filtering was performed. To calculate PC importance, scree plots were generated in Python by computing the variance explained by each PC and then plotting. For the full scripts please see Github (https://github.com/pilaralv/GBA1-multiomics).

To calculate if mutation severity was correlated to changes in gene expression, we ran linear regressions. To do this, we recoded each genotype as 1-4, (WT=1, heterozygous=2, homozygous=3, KO=4) under a new variable “GBA1_GROUP”, assuming a linear relationship between the four categories. Linear regressions were run in Python with these recoded groups against each gene for each cell type. Per assay, we included as covariates PCs that explained at least 80% of variance. For RNA, we corrected for 2 PCs for all cell types. Multiple test correction was performed using Benjamini-Hochberg method. After correction, significant hits were determined as those with an FDR < 0.05.

### Metabolomics data processing and analysis

To generate a cleaned data table, we took the batch normalized data received from Metabolon, and mapped the metabolite ID to the metabolite name based on the chemical annotation provided. Then, we edited the metabolite names to remove any special characters so that they would be properly read in downstream analyses. Additionally, we added an “X_” prefix to each metabolite to avoid issues with number-starting names. ata was not merged across cells as it was generated in two batches.

PCs were calculated using prcomp and plotted with autoplot in R for each cell type. We removed any metabolite where there was no variance across all entries as those interfered with the PCA. No additional filtering was performed. To calculate PC importance, scree plots were generated in Python as described above.

For the linear regressions, we recoded our variables as described above and corrected for 2 PCs for all cell types. Multiple test correction was performed using Benjamini-Hochberg method and significant hits were determined with the same threshold as above.

Additionally, we generated heat maps of regression results for the metabolites forming part of the ceramide subclass. These included: ceramide PE, ceramides, dihydroceramides, hexosylceramides, and lactosylceramides. Heatmaps were generated using ggplot2 in R. We also used the ggplot2 package to visualize regression results across each -omic.

### Somalogic proteomics data processing and analysis

The .adat file was manipulated in R to get a final table of our proteins of interest. First, we generated a file mapping the protein sequence ID to the protein name, and filtered out any non-human proteins. Then, we updated the data file by replacing the protein sequence ID with the corresponding protein name. This same process was done to generate matrices for all three cell types. Due to low Somalogic QC, we removed four neuronal outliers (WT_rep3, D448H_homozygous x 3). Additionally, as the data was generated in two batches it was not possible to merge across the three cell types.

PCs were calculated using prcomp and plotted with autoplot in R for each cell type. We removed any protein where there was no variance across all entries as those interfered with the PCA. No additional filtering was performed. To calculate PC importance, scree plots were generated in Python as described above.

For the linear regressions, we recoded our variables as described above and corrected for 2 PCs for the IPSC, and 3 PCs for the forebrain cortical neurons and microglia. Multiple test correction and threshold determination was done as described above.

## Results

### Characterization and validation of the *GBA1* KOLF2.1J isogenic model

After CRISPR editing, we obtained 3 clones of 5 edited KOLF2.1J cell lines including *GBA1* p.D409H, *GBA1* p.D409V (both in heterozygous and homozygous state), and the *GBA1* knockout, as well as the KOLF2.1J parental line (WT). The two variants p.D409H (also known as p.D448H, rs1064651) and p.D409V (also known as p.D448V, rs77369218) are known to be pathogenic. Before performing differentiation protocols and experiments, we validated the isogenic model extensively and completed multiple quality control steps at the RNA and protein level.

iPSC cells were cultured and RNA and protein were isolated from cell pellets. We performed Illumina RNA sequencing to assess *GBA1* transcription and to confirm the engineered variants. Variant calling of the RNA sequencing data for each line confirmed that the variants of interest were present. Additionally, the mapped RNA sequencing files were manually verified on IGV to confirm the presence of each variant (Supplementary Figure 1). Importantly, total *GBA1* RNA was significantly reduced in the knockout line (P= 1.96x10^-6^, β=-1.36, β_SE=0.175) (Figure 1A); however, there were still some sequence reads appearing to map to *GBA1* in the RNA data. This is most likely due to multi mapping of reads from *GBA1LP*, a highly homologous pseudogene (Supplementary Figure 2).

**Figure 1:**
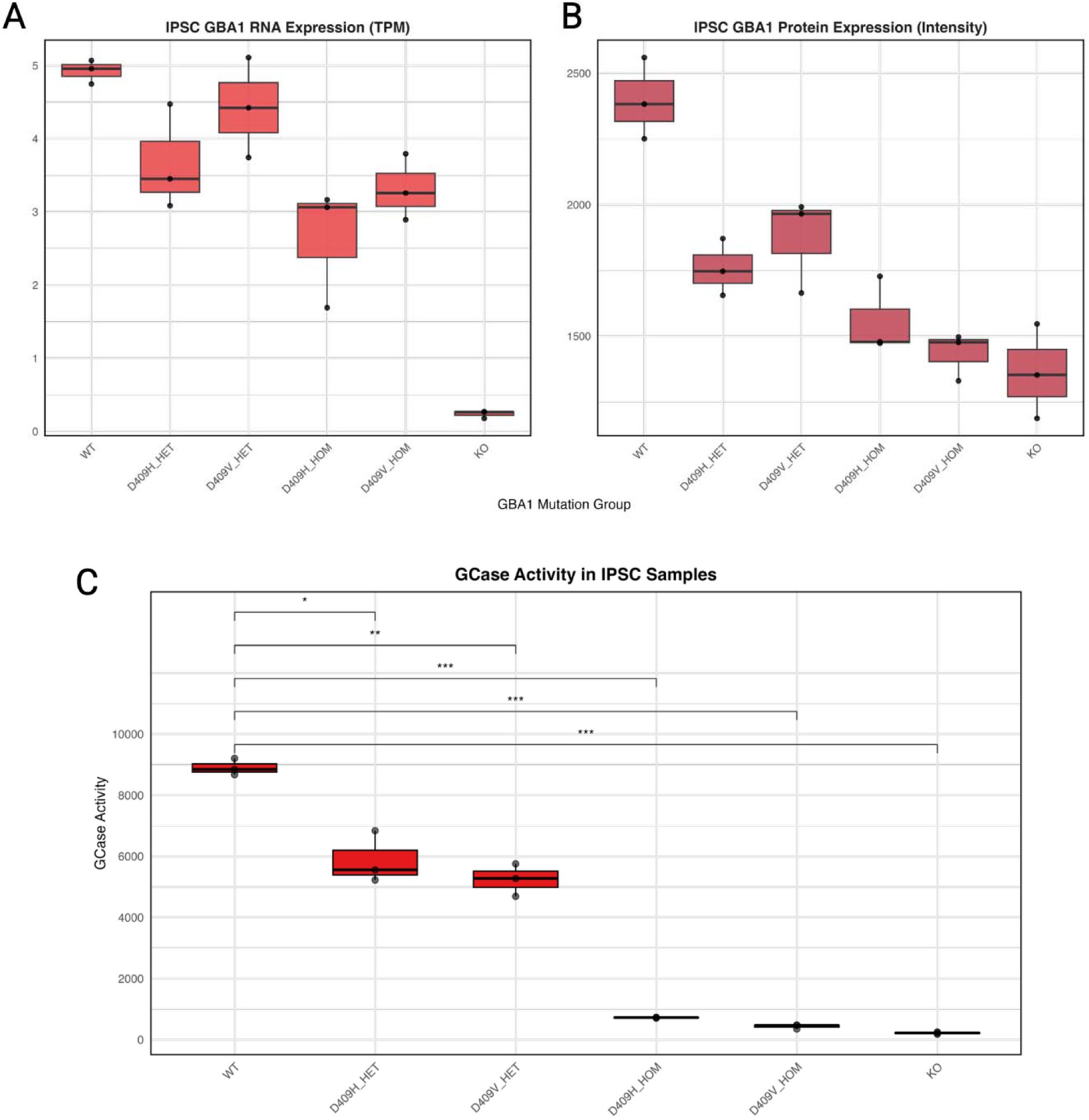
iPSC Illumina RNA sequencing, SomaLogic proteomics, and GCase activity across *GBA1* genotypes. A) RNA sequencing of the *GBA1* allelic series shows some reduction in *GBA1* expression and, importantly, almost abolishes *GBA1* expression in the *GBA1* knockout (KO), with some expression left due to mismapped sequence reads. B) SomaLogic proteomics data showed a consistent reduction of GCase protein level in the allelic series. C) Similarly, when measuring GCase activity, enzymatic activity is reduced in the allelic series. Stars represent significance in a t-test of genotype group vs the wild type group. One star represents p < 0.05, two stars represent p < 0.005, and three stars represent p < 0.0005. For all panels, the middle boxplot line represents the second data point and the box outline represents the first and third quartiles.

Next, we explored proteomic data generated using Somalogic SomaScan Assay 7K to investigate the effects of the engineered variants on GCase protein levels. This data showed that both heterozygous and homozygous variant lines, and the *GBA1* knockout line, had significantly reduced GCase protein levels compared to the wild type (P=4.50x10^-6^, β=-317.13, β_SE=43.99) (Figure 1B).

Finally, we confirmed that there is reduced enzymatic activity in the single variant *GBA1* cell lines and knockout *GBA1* lines, which highly correlated with the proteomic expression results (Figure 1C), indicating its utility for further experiments. Statistical analysis revealed that all genotype groups were significant when compared against the wild type group at a p< 0.05. The order of significance across groups from most significant to least significant is as follows: D409V_HOM-WT (p=1.50 x 10^-4^, β = 8472.80, β_SE = 162.11), KO-WT (p=2.61x10^-4^, β = 8689.13, β_SE=158.19), D409H_HOM-WT (p=3.56x10^-4^, β= 8186.38, β_SE=156.99), D409V_HET-WT (p=1.84x10^-3^, β= 3667.63, β_SE= 345.06), D409H_HET-WT (p= 0.018, β=3033.64, β_SE=516.05).

### Differentiation of isogenic *GBA1* iPSCs to forebrain cortical neuronal and microglia cell types

After characterizing iPSC edited lines, we expanded the isogenic model into additional brain-relevant cell types, namely forebrain cortical neurons and microglia. To do so, we differentiated the 18 KOLF2.1J derived lines (6 genotypes at 3 replicates/clones) into cortical neurons and microglia for a total of 36 lines (Figure 2). Differentiated cells were harvested after 25 days (for neurons) and 30 days (for microglia) and underwent the same data generation as the iPSC lines (RNAseq, proteomics, GCase activity for neurons only) and in addition, metabolomic assessments were performed on all lines and cell types.

**Figure 2:**
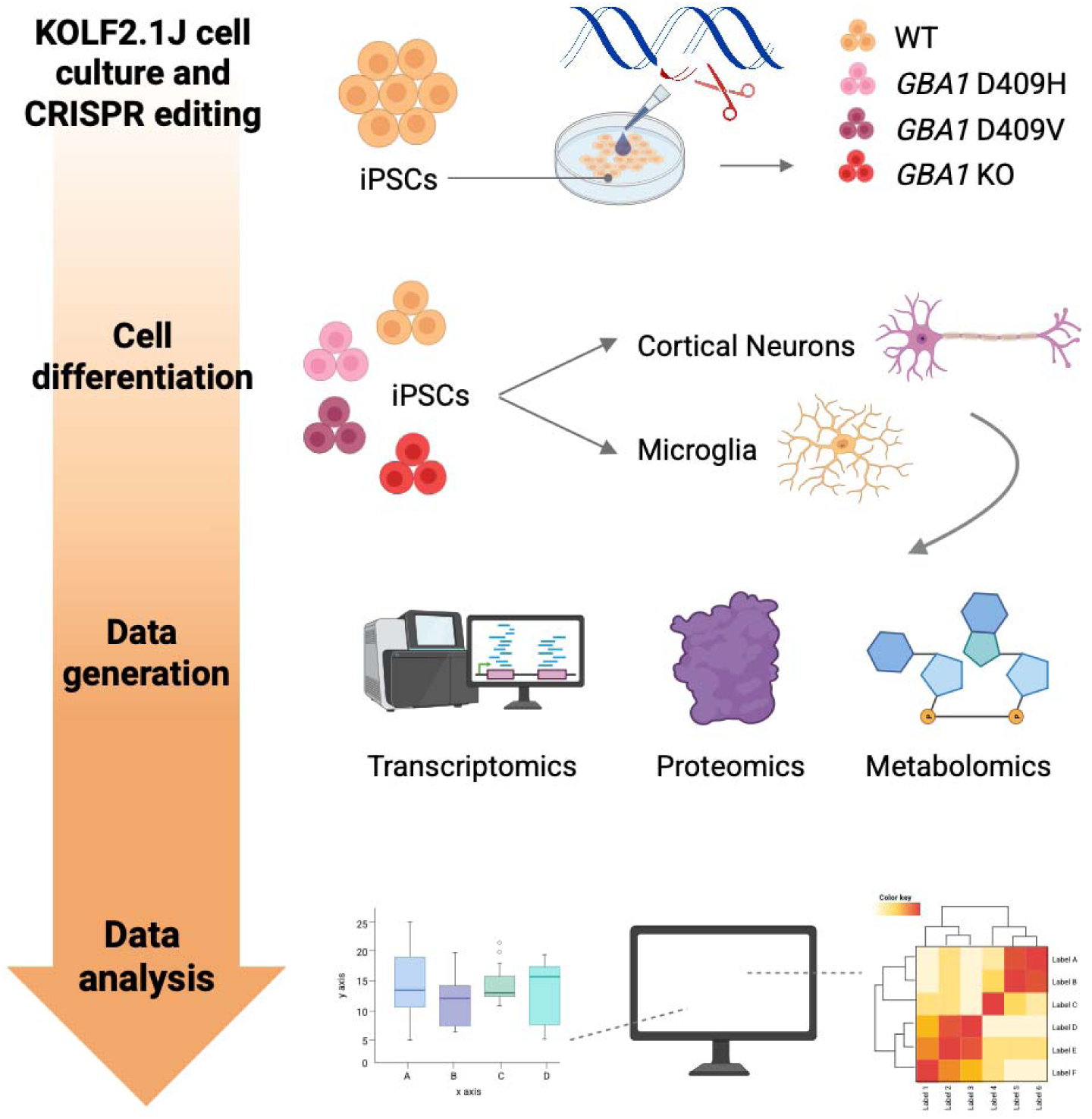
Graphical representation of the experimental design. KOLF2.1J wild type (WT) iPSCs underwent CRISPR editing for *GBA1* mutations. *GBA1* p.D409H and p.D409V edits were performed to generate both heterozygous and homozygous lines. Each edited line was then differentiated into forebrain cortical neurons and microglia, with three clones each for 18 samples per cell type and 54 lines total. These samples then went through transcriptomic, proteomics, and metabolomics data generation and the data were analyzed to look for potential mechanisms in across data modalities and cell types.

We examined gene expression levels of cell-type marker genes, *POU5F1*, *MAP2*, and *TREM2,* to label iPSC, neurons, and microglia, respectively (Figure 3A). Similarly, we assessed the protein expression data of marker genes SOX2, DPYSL5 and TREM2 in iPSC, neurons, and microglia, respectively (Figure 3B). The expressions of the marker genes matched expected values, with high expression in the appropriate cell type and limited to no expression in the other cell types. This gives confidence that the cell-type differentiations were successful and thus increases reliance on downstream cell-specific results. As done previously for the iPSC allelic series, we wanted to confirm the predicted GCase activity reduction by genotype in our neuronal lines. The GCase results show that activity decreases corresponding to the severity of the genotype for the forebrain cortical neurons, following the trend seen in iPSCs, although levels varied across cell types (Supplementary Figure 3).

**Figure 3:**
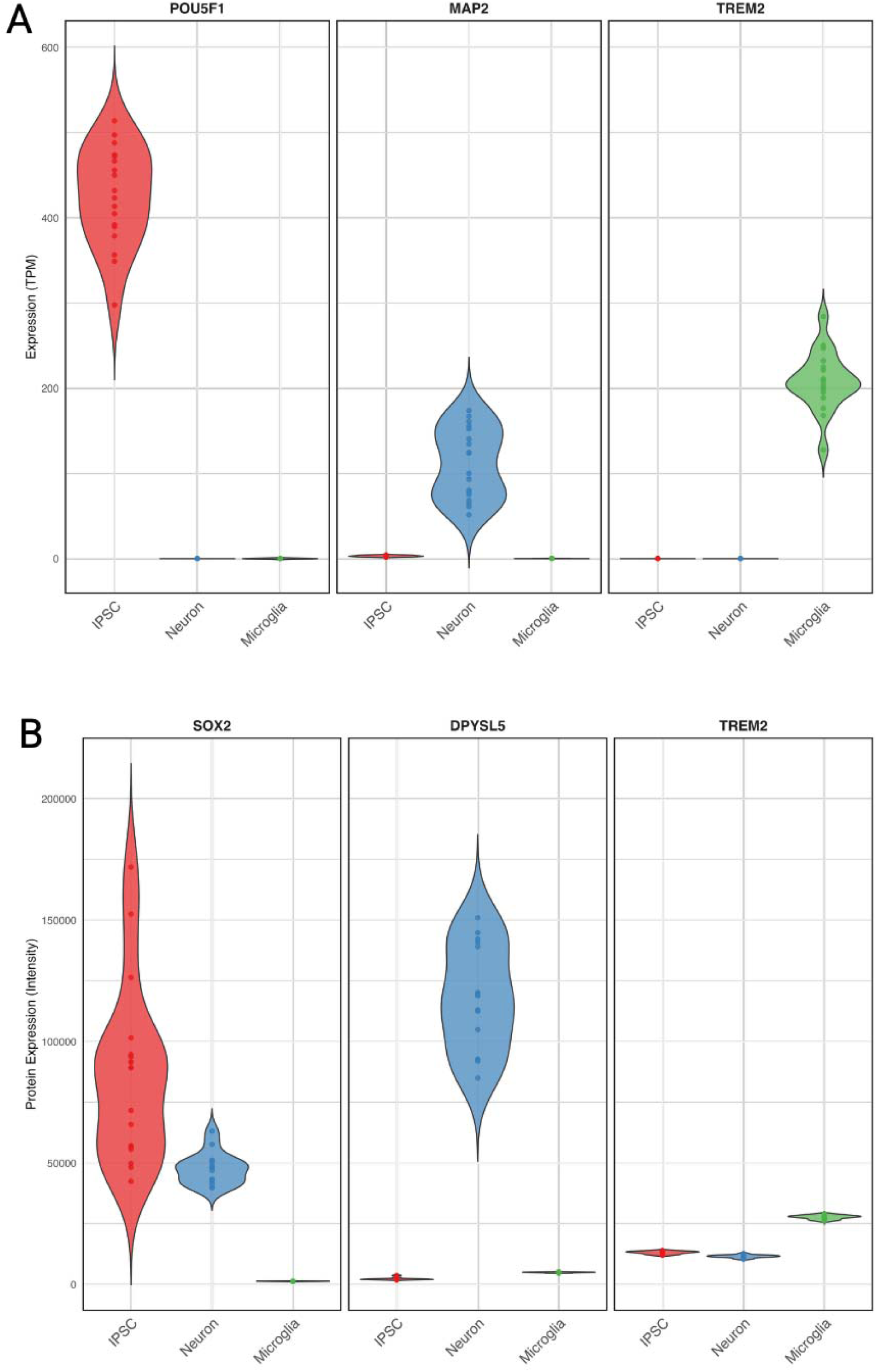
Marker gene analysis based on RNA sequencing and proteomics data across cell types. A) RNA sequencing reveals apparent differences across cells for each respective marker gene thus indicating successful differentiation. *POU5F1* is a key transcription factor for maintaining the pluripotency and self-renewal of IPSCs. *MAP2* encodes a protein that stabilizes microtubules, playing a crucial role in neuronal development and is highly enriched in the dendrites of mature neurons. Lastly, *TREM2* encodes a transmembrane receptor primarily expressed in microglia and plays a role in microglial activation. B) Proteomic data confirms successful differentiation as each cell type significantly expresses respective marker proteins. SOX2, like POU5F1, is a transcription factor essential for maintaining pluripotency in IPSCs. DPYSL5 is vital for neuronal differentiation and axon guidance.

### Multi-omics analysis of differentiated cell types

Next we evaluated the RNA, protein, and metabolomic datasets. For all assays, principal component analysis (PCA) plots were generated and regressions were run with genotypes recoded as 1-4 (WT=1, heterozygous=2, homozygous=3, KO=4). We will describe each -omic analysis below.

#### Illumina RNA sequencing across *GBA1* genotypes

After sequence data processing, we calculated expression for 60,880 gene entries (protein coding and non protein coding) from hg38. iPSC, neuron, and microglia tables were merged and any entry with an average TPM below 2 across all 36 samples was removed. After filtering based on expression, 12,205 genes remained, which were then used to calculate principal components across the three cell types. PCA revealed that the main clustering came from cell type as opposed to genotype (Figure 4A). Notably, the microglia were the most tightly clustered overall and the forebrain cortical neurons the least clustered, potentially due to a less homogeneous differentiation protocol for the neurons. When making the individual tables with principal components for the downstream regression, we were left with 12,109 genes for iPSC, 11,491 for neurons, and 9,492 for the microglia.

**Figure 4:**
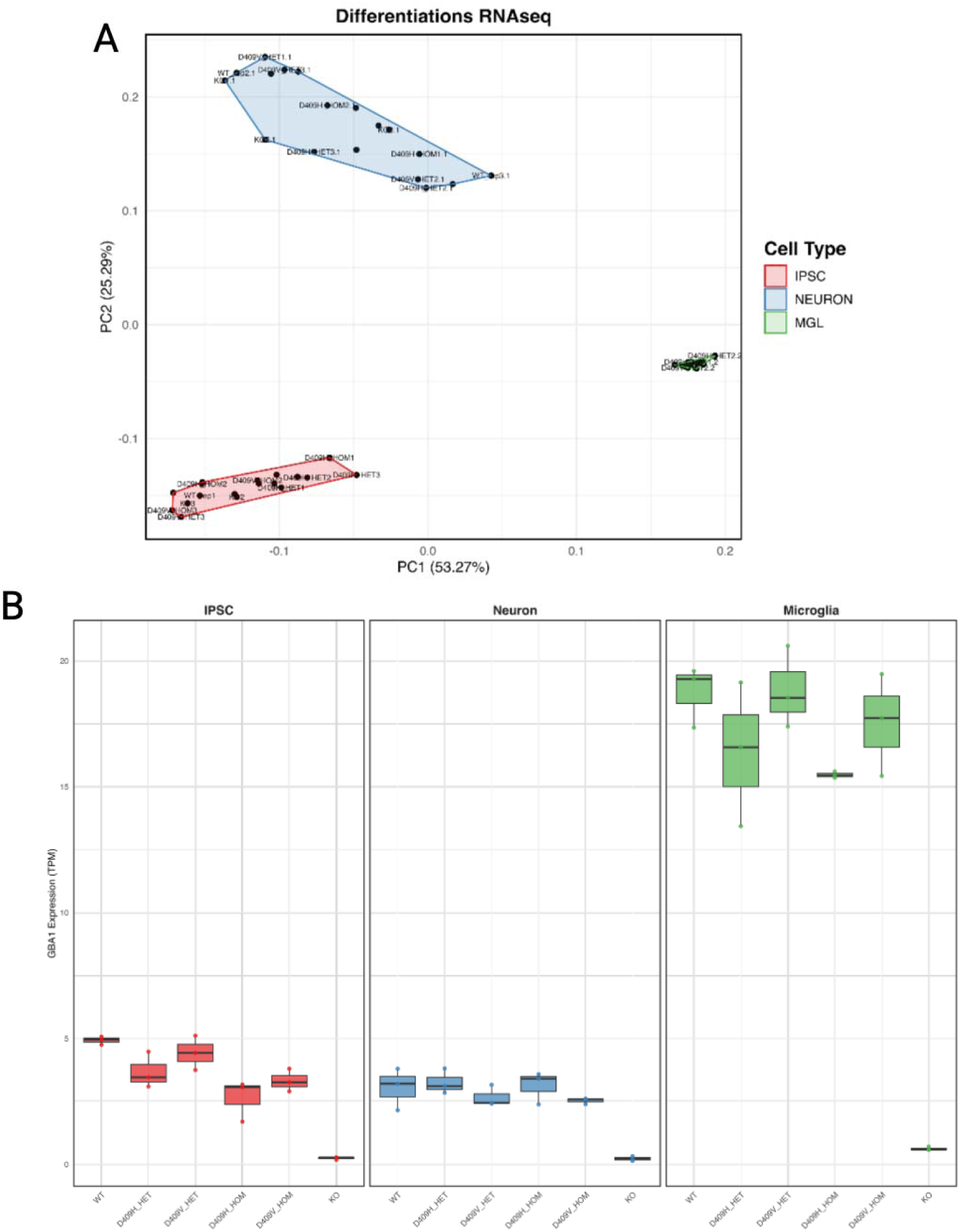
Transcriptomic differences across cells and genotypes. A) Principal component analysis of transcriptomic data shows clustering by cell type rather than genotype. B) Illumina RNAseq reveals a reduction in *GBA1* expression with increasing severity of genotype, particularly in the knockout lines. For all panels, the middle boxplot line represents the second data point and the box outline represents the first and third quartiles.

At the RNA level, we observed a reduction in *GBA1* expression as the genotype severity increases across all three cell types. This was statistically significant in all cell types, however for both neurons and microglia the significance was lost after multiple test correction for all genes tested, (IPS_FDR=0.024, IPS_p= 1.96x10^-6^, IPS_β=-1.36, IPS_ β_SE=0.175, Neuron_FDR=0.281, Neuron_p= 4.55x10^-3^, Neuron_β=-0.841, Neuron_ β_SE=0.249, MGL_FDR=0.076, MGL_p=3.92x10^-3^, MGL_β=-3.24, MGL_ β_SE=0.939). While the microglial lines had *GBA1* expression levels that were up to 4 times higher than the other two cell types, the reduction by genotype was most notable in the undifferentiated iPSCs. Of note, we did not see this reduction in *GBA1LP*, the pseudogene, which seems to be unaffected by the introduction of the pathogenic variants in the KOLF2.1J cell line, nor did we see any correlation between *GBA1* and *GBA1LP* expression (Supplemental Figure 4A-B).

Additionally, there were significant changes in the expression levels of two other genes in iPSCs. *FREM2* (FDR=0.032, p=6.21x10^-6^, β=-1.09, β_SE=0.156), which provides instructions for an extracellular matrix protein and mutations in this gene case Fraser syndrome (OMIM #617666), and *HEPH* (FDR=0.032, p=7.84x10^-6^, β=-1.78, β_SE=0.260), a gene encoding the hephaestin protein involved in iron and transport (Supplemental Figure 5). In the forebrain cortical neurons, two other genes were statistically significant, *ZNF484* (FDR=0.021, p=1.81x10^-6^, β=-0.336, β_SE=0.043), in the zinc finger protein family and *SH3PXD2B* (FDR=0.021, p=3.69x10^-6^, β=0.672, β_SE=0.092) encoding a protein involved in cell adhesion and migration Supplementary Figure 6. Lastly, there were 179 genes in the MGL that were statistically significant. Notably, none of these overlapped with the iPSC or neuronal hits, suggesting cell-type-specific differences across how cells types respond to *GBA1* damaging variants. For the full regression tables and statistics, see Supplementary Tables 3-5.

#### SomaLogic proteomics across *GBA1* genotypes

Proteomic data was run through SomaLogic on two batches, the first included the iPSCs and neurons, and the second the microglia. Because of the nature of the assay which measures intensities rather than absolute values at each batch collection, we could not merge the data for the PCA. As such, we generated the PCA plots for the iPSC and neurons together and then plotted the microglia separately (Supplemental Figure 7A-B). Due to quality control issues, we had to remove four neuronal samples before running our analysis (WT_rep3, D409H_homozygous x 3 all marked as significant outliers). For the first batch, which included the iPSC and neurons, we identified 7335 proteins across both. For the microglia in the second batch, we identified 7339 proteins. As with the RNA, the data clustered more by cell type but with no obvious clustering by genotype in the PCA. After multiple test corrections, only the GCase protein (encoded by *GBA1*) was statistically significant for all three cell types (IPS_FDR=0.033, IPS_p=4.5 x10^-6^, IPS_ β=-317.13, IPS_ β_SE=43.99, Neuron_FDR=0.002, Neuron_p=2.23x10^-7^, Neuron_ β=-336.86, Neuron_β_SE=27.28, MGL_FDR=1.97x10^-5^, MGL_p=2.68x10^-9^, MGL_ β=-1905.57, MGL_ β_SE= 134.07). No other protein surpassed the threshold. We also inspected the LIMP2 (encoded by *SCARB2*) protein, which is a known lysosomal membrane protein. However, this had both FDR and p values > 0.05 across all three cell states (p=0.843/FDR=0.916 in iPSCs, p=0.045/FDR=0.855 in neurons, p=0.707/FDR=0.993 in microglia). For the full regression tables and statistics, see Supplementary Tables 6-8.

**Figure 5:**
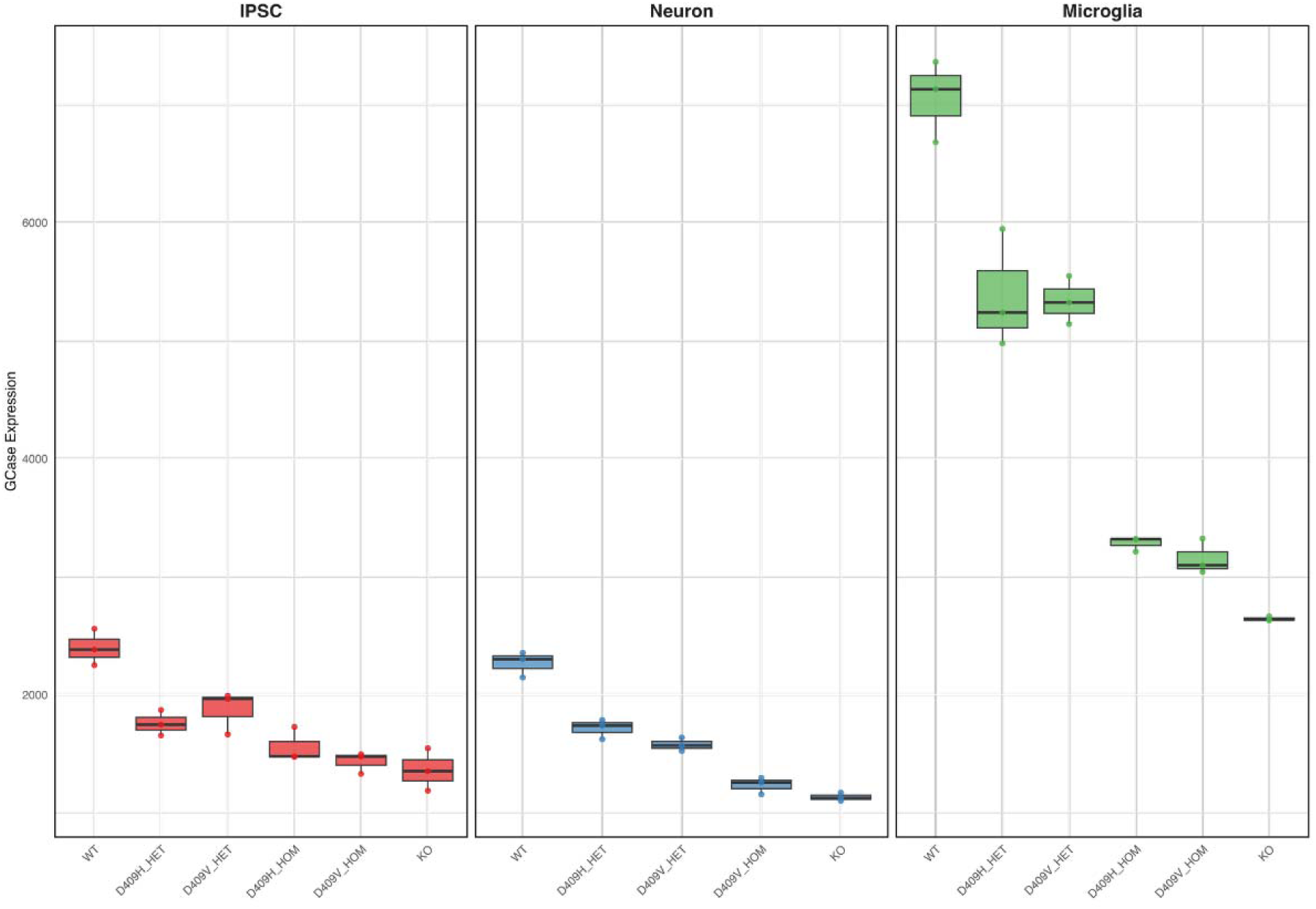
Proteomics differences across cells and genotypes. A) SomaLogic proteomics reveals a reduction in GCase protein expression with increasing genotype severity, particularly in the knockout lines. For all panels, the middle boxplot line represents the second data point and the box outline represents the first and third quartiles.

#### Metabolon metabolomics across *GBA1* genotypes

Like the proteomic data above, the metabolomics data was run in two batches, with one including the iPSC and neurons, and the second the microglia. Due to the nature of the metabolomics assay, different metabolites can be detected each time it is run; thus, the data cannot be merged across batches. To calculate principal components, we filtered out metabolites whose values did not vary across all samples. We identified 636 metabolites for batch one and 710 metabolites for batch two. When making the individual tables for the regression analysis, we were left with 606 metabolites for iPSCs, 534 for neurons, and 710 for microglia. Each of the metabolites is classified under a “super_pathway” and a “sub_pathway” according to Metabolon. Of note, some super pathways include the terms: peptide, amino acid, carbohydrate, lipid, etc, while the sub pathways involve terms: ceramides, aminosugar metabolism, dipeptide, lypophospholipid, etc. For a full key of metabolites and their pathways see Supplementary Table 9. While nothing was significant at an FDR < 0.05, we had a high proportion of metabolites that belonged to the ceramide sub-pathways as top hits, especially in the neurons (p < 0.05). These differences were primarily seen in the homozygous and knockout lines (D409H/V homozygous and KO). As such, we extracted out any metabolite belonging to these sub-pathways and generated heat maps across all three cell types (Figure 6). Specifically, these pathways included ceramide PE, ceramides, dihydroceramides, hexosylceramides, and lactosylceramides, encompassing a total of 30 metabolites. Naturally, the ceramide pathway is highly interesting given that GCase is crucial for sphingolipid metabolism, breaking down glucosylceramide into ceramide and glucose. Some examples of interest include glycosylceramide (Supplemental Figure 8A) and glycosyl-N-nervonoly-sphingosine (Supplemental Figure 8B). For the full regression tables and statistics, see supplementary tables 10-12. For ceramide levels plotted, see supplementary table 13.

**Figure 6:**
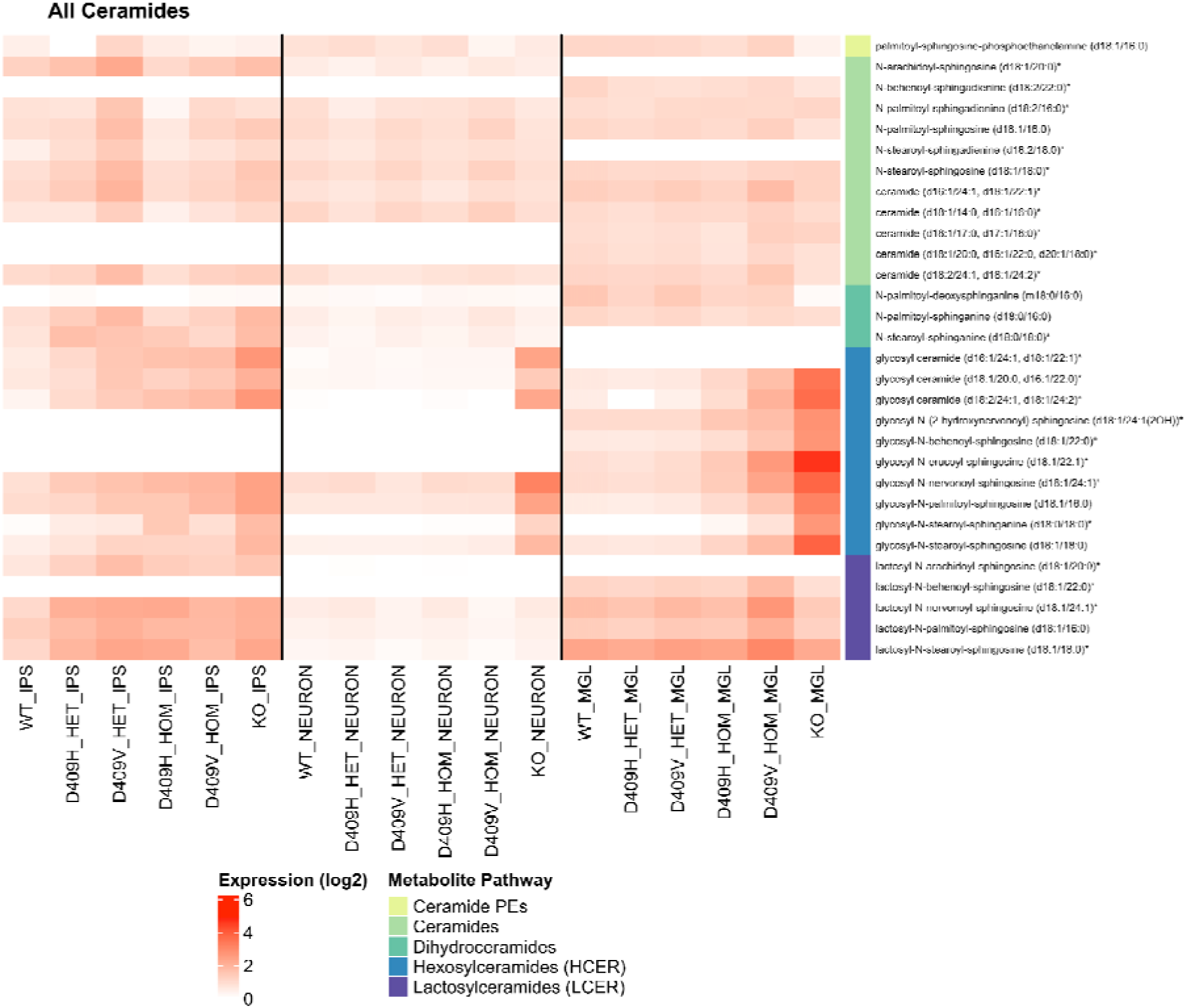
Heatmap of ceramide pathways across cells. Metabolomic analysis revealed that metabolites associated with ceramide pathways are differentially expressed across genotypes. Of note, there is an increase in levels of hexosylceramides in the microglial *GBA1* knockout lines. A total of 30 metabolites were captured across two batches (IPSC and forebrain cortical neuron samples in the first batch and microglia samples in the second). Note that white lines include no values obtained from the metabolomics experiment. Original ceramide levels can be found in Supplementary Table 13.

## Discussion

This study aimed to characterize and validate an isogenic KOLF2.1J iPSC model harboring *GBA1* variants associated with GD, LBD and PD. We successfully validated a comprehensive allelic series generated through CRISPR/Cas9 editing, including heterozygous and homozygous p.D409H and p.D409V variants, as well as a *GBA1* knockout. Extensive characterization at the RNA, protein, and enzymatic activity levels confirmed the expected reduction of GCase expression and activity with increasing genotype severity in iPSCs.

In addition, we describe here a pilot experiment to dissect the effects of *GBA1* mutations in an isogenic model using a multi-omic approach. The KOLF2.1J cell line, a suggested standard for neurodegenerative studies, was edited to harbor *GBA1* variants and was knocked out across three cell states: iPSC, neurons, and microglia. Transcriptomic, proteomic, and metabolomic data were generated for all samples to investigate potential additional targets in *GBA1* based disease mechanisms. By using an isogenic model, we are able to control variability stemming from a difference in genetic background across donors. iPSCs were successfully differentiated into brain-relevant cell types (forebrain cortical neurons and microglia), as evidenced by high expression of the marker genes for each cell type. Of interest is the high *GBA1* expression in microglia which is an understudied cell type in the context of *GBA1*. Importantly, both transcriptomic and proteomic data showed reduced RNA expression and protein abundance for *GBA1* with increasing *GBA1* variant severity across all cell types This data also aligns with the GCase assay data showing reduced activity with *GBA1* variants of increased severity.

While our statistical analyses did not reveal clear targets after multiple test correction, using this method in larger-scale studies may elucidate cofounding factors impacting *GBA1* disease mechanisms and potentially penetrance. Interestingly there was very limited overlap between RNA and protein results across cell types, which raised the question if different cell types are differently affected by *GBA1* pathogenic variants. Additionally, our metabolomic data points to alterations in ceramide pathways with *GBA1* mutations as a mechanism of interest which is in line with the current literature around GBA1 and thus tells us that perhaps metabolites should be increasingly prioritised in *GBA1* or lysosomal risk mechanisms over RNA based methods.

This study has multiple limitations. Due to our small sample size, and its pilot design, it is difficult to achieve statistically significant results with robust methods correcting for principal components and multiple test corrections. Large scale differentiation of isogenic cell lines is costly and time-consuming, however with the development of automated cell culture robotics, it can now be performed in certain laboratories. For example, there are currently efforts led by iNDI at the National Institutes of Health using high throughput robots to grow and differentiate cells so that future isogenic experiments can be run with larger numbers of samples. Additionally, for the metabolomic data, glucosyl- and galactosyl-ceramides were measured together as a single readout. This could limit the changes observed with *GBA1* variants, particularly in cell types (i.e. neurons) where galactosyl species are more abundant than glucosyl species. Additionally, despite using a multi-omic approach, we needed to look at each - omic read-out individually, as tools are not available to seamlessly integrate different data output (genes, proteins, metabolites, enzymatic activity). As multi-omic machine learning approaches improve, they also need a larger sample size and future larger studies should focus on integrating all of the generated data to create a network which may elucidate significant hits and pathways missed during this preliminary analysis.

Our study successfully validated an isogenic KOLF2.1J iPSC model carrying pathogenic *GBA1* variants. This model effectively replicates the anticipated decrease in glucocerebrosidase (GCase) expression and activity across various cell types and uncovers significant disruptions in ceramide metabolism. Our initial multi-omics analysis delivers a thorough perspective on the potential molecular impacts of altered GCase, underscoring the power of this model to enhance our understanding of *GBA1*-associated diseases, particularly when expanded to include a wider range of *GBA1* variants and cell types. Furthermore, our findings suggest that proteomic and metabolomic analyses may offer more valuable insights than RNA-based studies into the effects of *GBA1* variants. Importantly, all data and code from this research are publicly accessible, paving the way for future investigations utilizing isogenic models to further clarify the pathogenic mechanisms of *GBA1*-related diseases and to facilitate the development of innovative therapeutic approaches. Future research should prioritize validating these observations in larger studies and with a broader spectrum of *GBA1* variants to gain a deeper understanding of their downstream consequences.

## Supplementary Figures

**Supplementary Figure 1: Confirmation of successful CRISPR edits in KOLF IPSCs.** IGV screenshot of IPSC lines over exon 9 of *GBA1* shows successful editing of heterozygous and homozygous variants. Reduced sequence read coverage of *GBA1* over knockout lines compared to wild-type lines confirms successful knockout edit. Remaining coverage over the *GBA1* knockout lines are due to multimapping sequence reads with highly homologous pseudogene *GBA1LP*.

**Supplementary Figure 2: Coverage reduction over the whole of *GBA1* in the knockout lines compared to the wild type.** Total RNA was significantly reduced for the knockout lines and sequence reads mapping to *GBA1* from the RNA data is likely from multi mapping from reads coming from *GBA1LP*, a highly homologous pseudogene. There is normal coverage over exons as expected for the wild-type samples.

**Supplementary Figure 3: GCase activity in forebrain cortical neuronal samples decreases with damaging genotype.** Measured glucocerebrosidase enzyme activity decreases with increasing severity of genotype. As expected, the *GBA1* knockout line has essentially no enzymatic activity and this activity is also greatly reduced for the homozygous lines. This follows the same pattern as the IPSCs (Figure 1C). Stars represent significance in a t-test of genotype group vs the wild type group. One star represents p < 0.05, two stars represent p < 0.005, and three stars represent p < 0.0005. The middle boxplot line represents the second data point and the box outline represents the first and third quartiles.

**Supplementary Figure 4: No significant differences in *GBA1LP* expression across genotypes. A)** While *GBA1* expression is reduced across genotypes, we do not see the same pattern in the pseudogene (*GBA1LP*) confirming that the pseudogene is likely not a cofounding factor in our data. **B)** No correlation appears when comparing *GBA1* and *GBA1LP* expression at the sample level across any cell type. The middle boxplot line represents the second data point and the box outline represents the first and third quartiles.

**Supplementary Figure 5: Significant hits in IPSC RNA.** In addition to *GBA1*, *FREM2* and *HEPH* were the other two hits statistically significant at an FDR < 0.05 (FDR=0.032 for both). *FREM2* provides instructions for protein making in the FRAS/FREM complex, and *HEPH* encodes a protein involved in iron metabolism and transport. The middle boxplot line represents the second data point and the box outline represents the first and third quartiles. The middle boxplot line represents the second data point and the box outline represents the first and third quartiles.

**Supplementary Figure 6: Significant hits in neuron RNA. From the transcriptomic analyses, SH3PXD2B and ZNF84 were significant at FDR < 0.05 (FDR=0.021 for both).** In the neurons, 2 genes were statistically significant. These were *ZNF484* (FDR=0.021) and *SH3PXD2B* (FDR=0.021). *H3PXD2B* encodes a protein involved in cell adhesion and *ZNF484* is in the zinc finger protein family. The middle boxplot line represents the second data point and the box outline represents the first and third quartiles.

**Supplementary Figure 7: Principal component analysis of SomaLogic data.** A) PCAs from the IPSC and neuron batch reveal clustering by cell type and little clustering by genotype. B) Microglia PCA also reveals no tight clustering by genotype. Since data was run in two batches, PCs from all three cell types were not run together.

**Supplementary Figure 8: Metabolomic data reveals changes in hexosylceramides.** A) Glycosyl ceramide intensity rises with increasing severity of genotype, particularly in microglia. B) Glysol-N-nervonoyl follows the same pattern with increase of intensity with more severe genotypes. The middle boxplot line represents the second data point and the box outline represents the first and third quartiles.

## Author contributions

Conceptualization: PAJ, PAWC, MRC, SB, CB

Experiments were performed by:

- Cell culture and differentiation: PAWC, DP, EL, SB
- Sequencing experiments: PAJ, CA, XR, CBP, KP, LM, KJB, MR
- GCase assay: LMG, YC, ES
- CRISPR editing: JAM, GN, WCK

Data was analyzed by: PAJ, PAWC, DP, LMG, MBM, MAN, ABS, CB

Writing (original draft): PAJ, PAWC, CA, MRC, SB, CB

Writing (review and editing): PAJ, PAWC, DP, LMG, CA, MBM, EL, YV, CBP, KP, LM, MAN, XR, ABS, KJB, JAM, GN, WCS, MR, ES, MRC, SB, CB

## Code Availability

All scripts and code for this project can be found at: https://github.com/pilaralv/GBA1-multiomics

## Data Availability

All data generated for this project is uploaded to the ASAP CRN cloud and will be available after internal QC Data is processed using the provided code.

## Funding agencies

This work was supported in part by the Intramural Research Program of the National Institutes of Health including: the Center for Alzheimer’s and Related Dementias, within the Intramural Research Program of the National Institute on Aging and the National Institute of Neurological Disorders and Stroke (project numbers ZIAAG000534 and ZIAAG000535).

This work utilized the computational resources of the NIH STRIDES Initiative (https://cloud.nih.gov) through the Other Transaction agreement - Azure: OT2OD032100, Google Cloud Platform: OT2OD027060, Amazon Web Services: OT2OD027852. This work utilized the computational resources of the NIH HPC Biowulf cluster (https://hpc.nih.gov).

## Acknowledgments

We thank the team at The Jackson Laboratory (JAX) for their support with iPSC editing. We thank Brian Sellers for his assistance with SomaLogic data and analysis. We also acknowledge Jon Brenton at the University College of London for providing and supporting the Salmon code used in this work. We thank the NIH HPC Biowulf cluster (https://hpc.nih.gov) for their computational resources.

