## Supplementary figures and images for "Dissecting the biological impact of *GBA1* mutations using multi-omics in an isogenic setting"

### Supplemental Figures

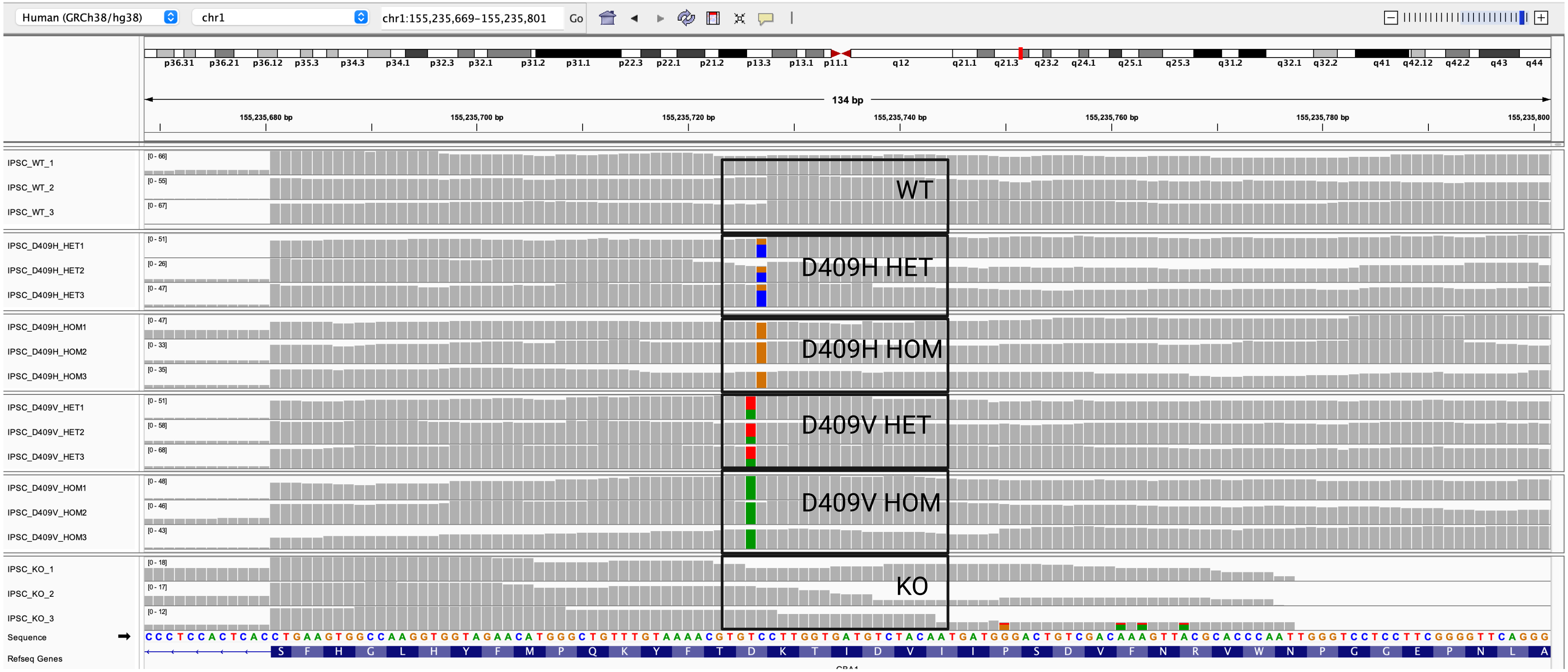

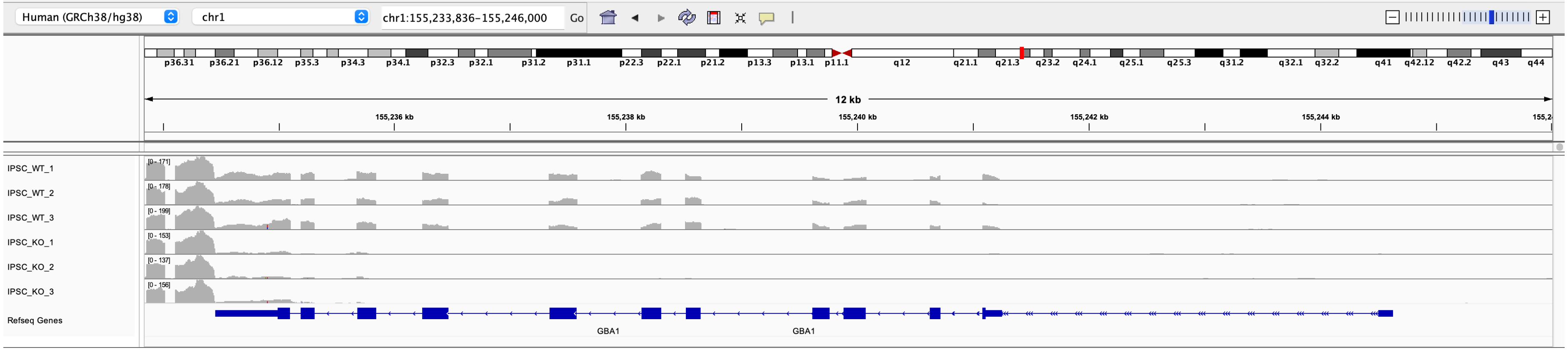

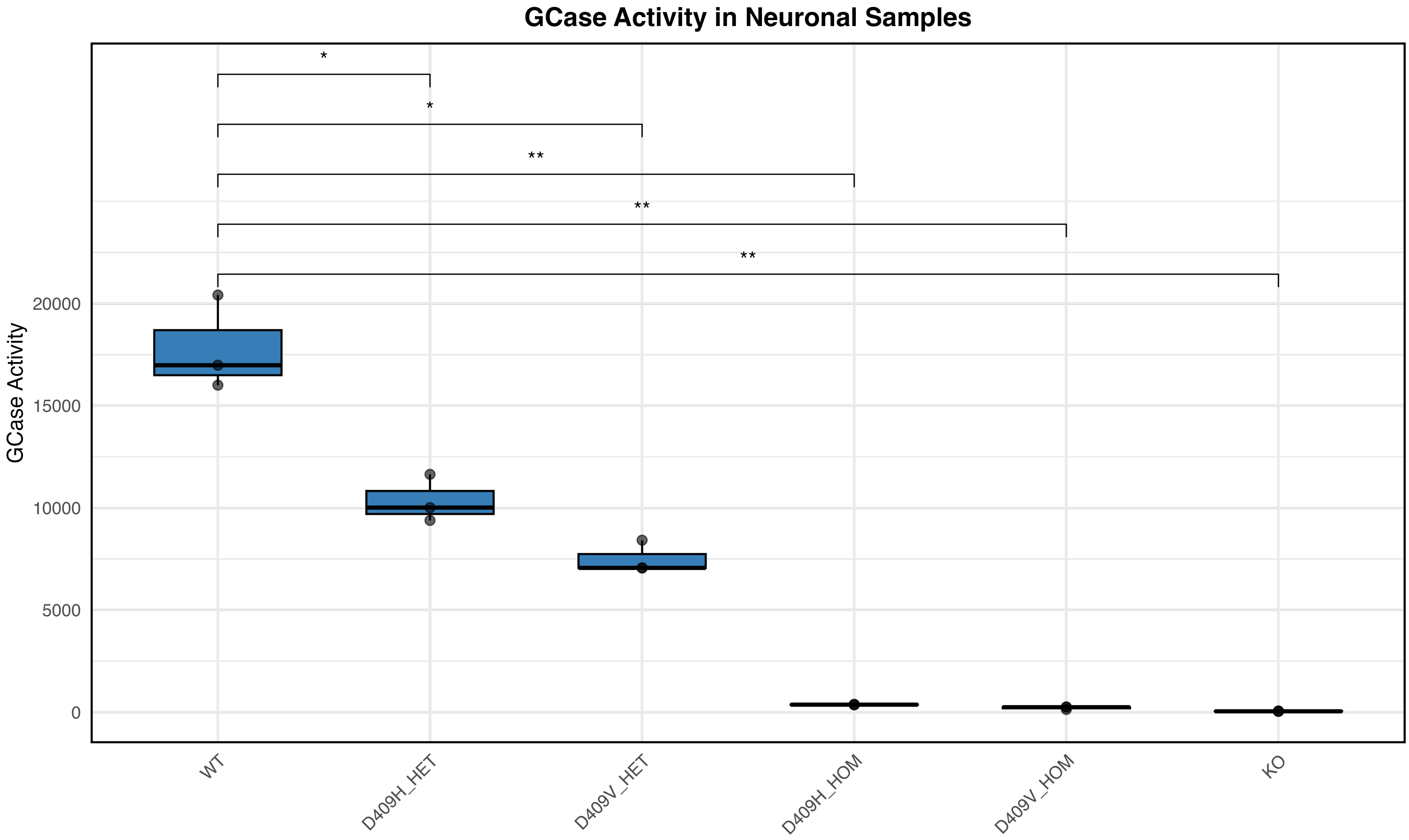

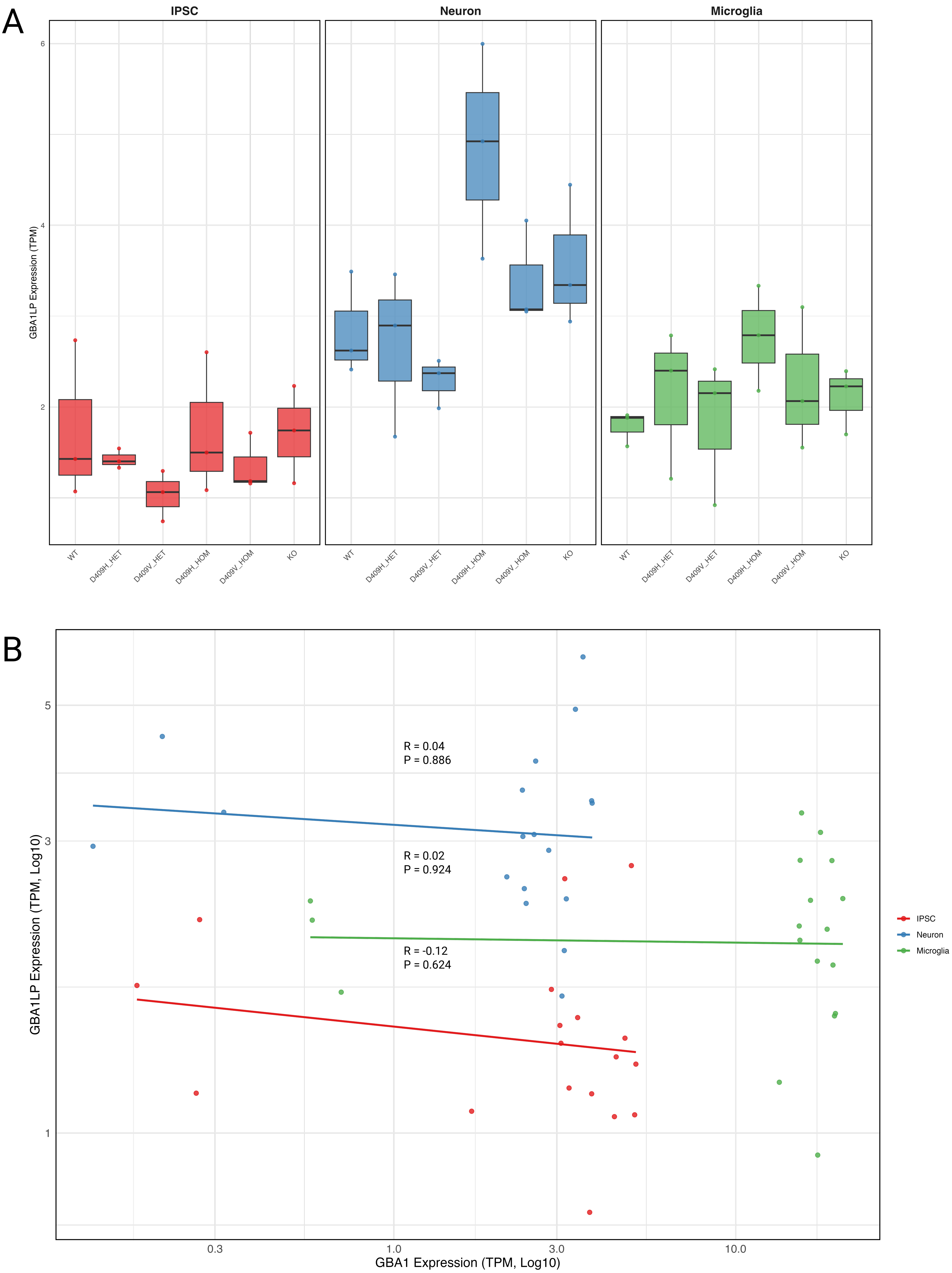

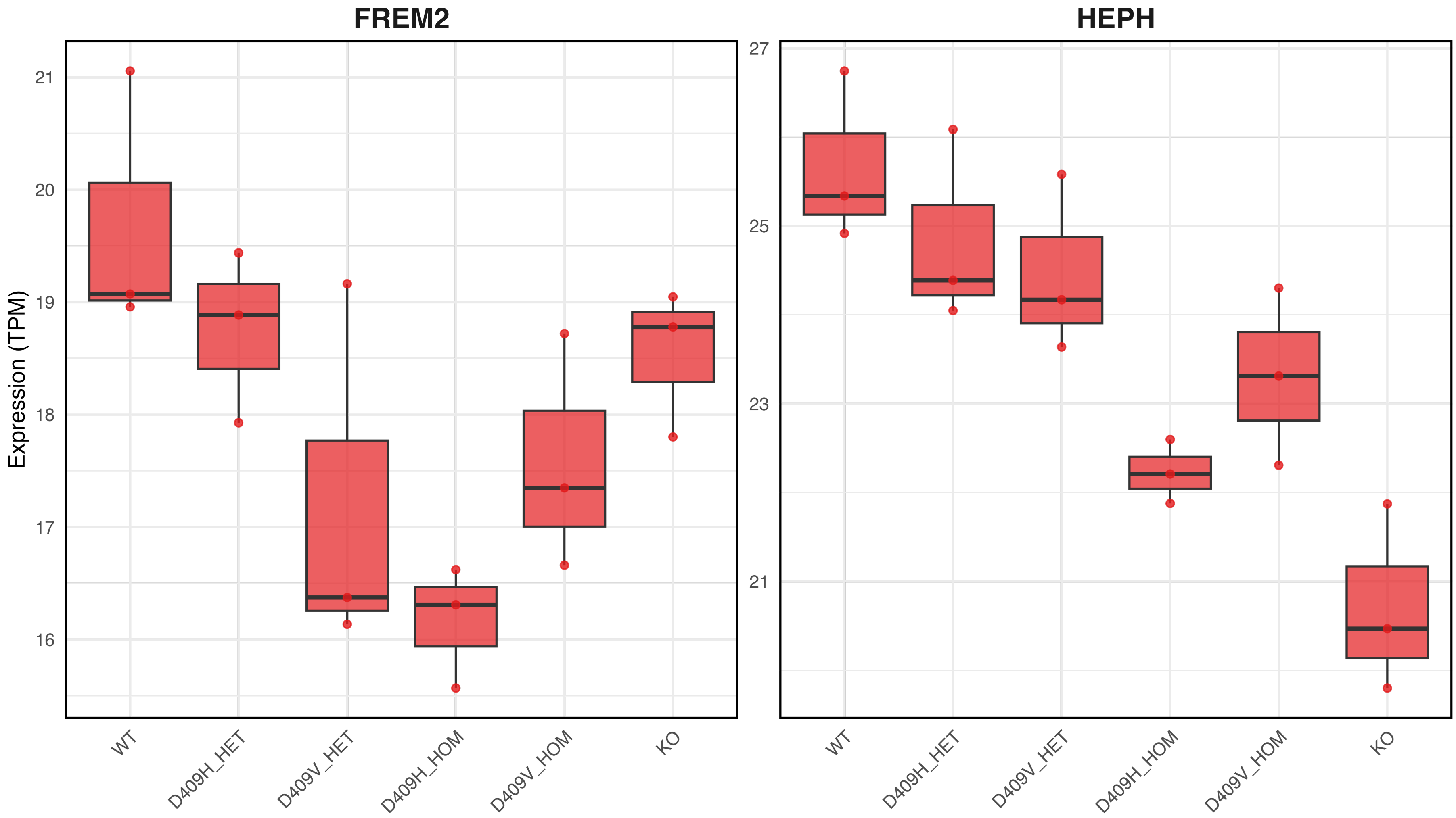

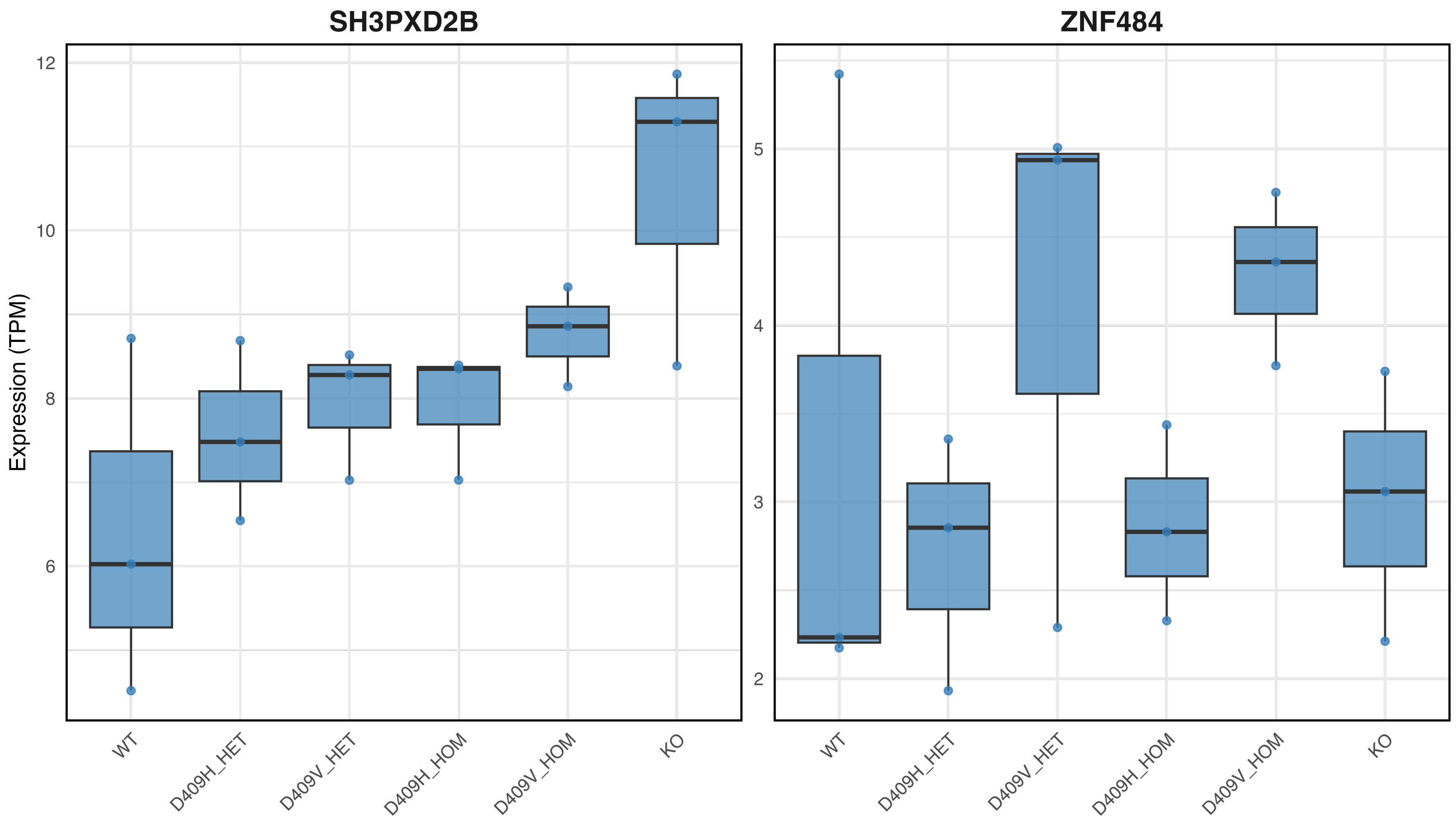

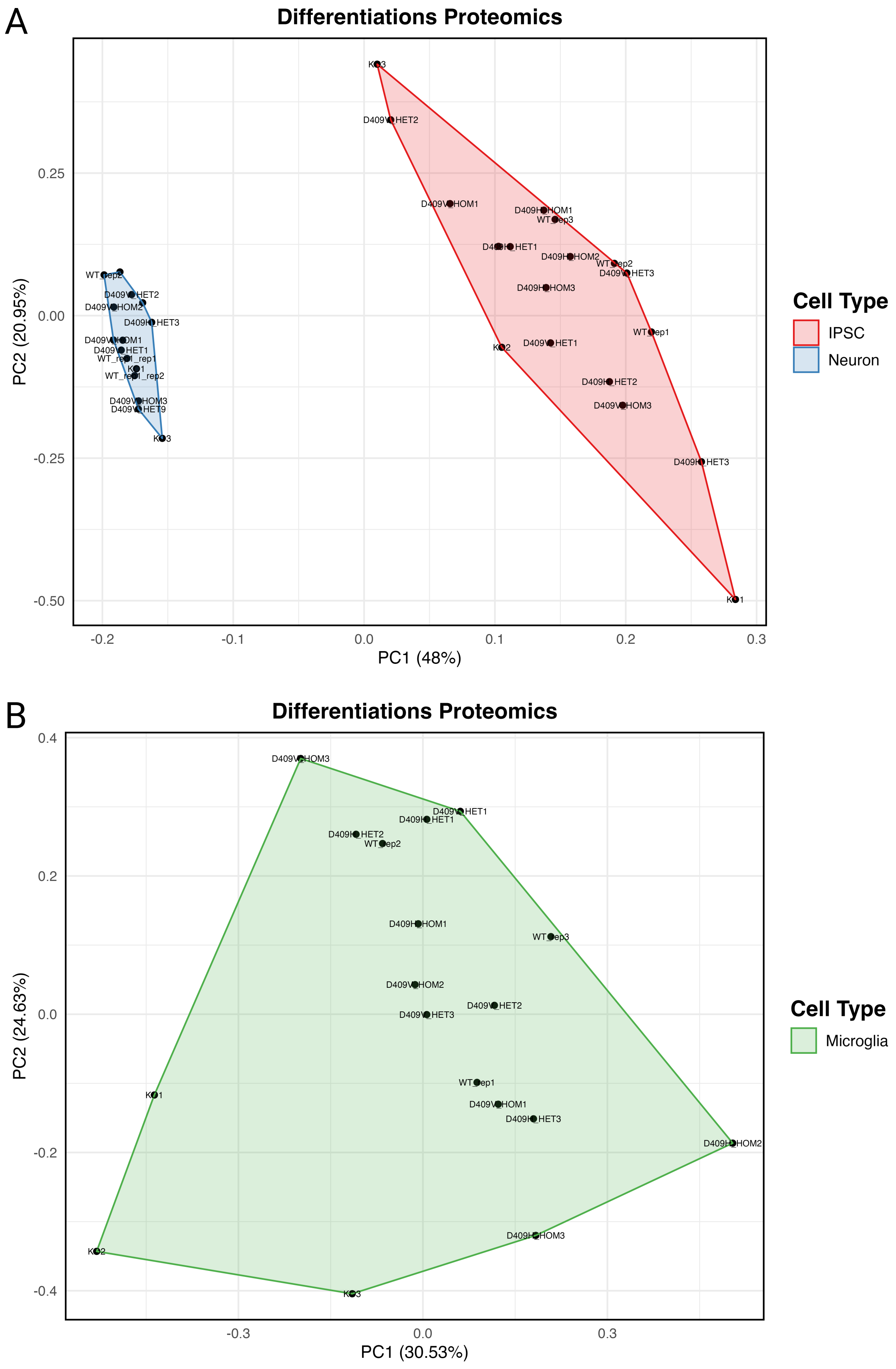

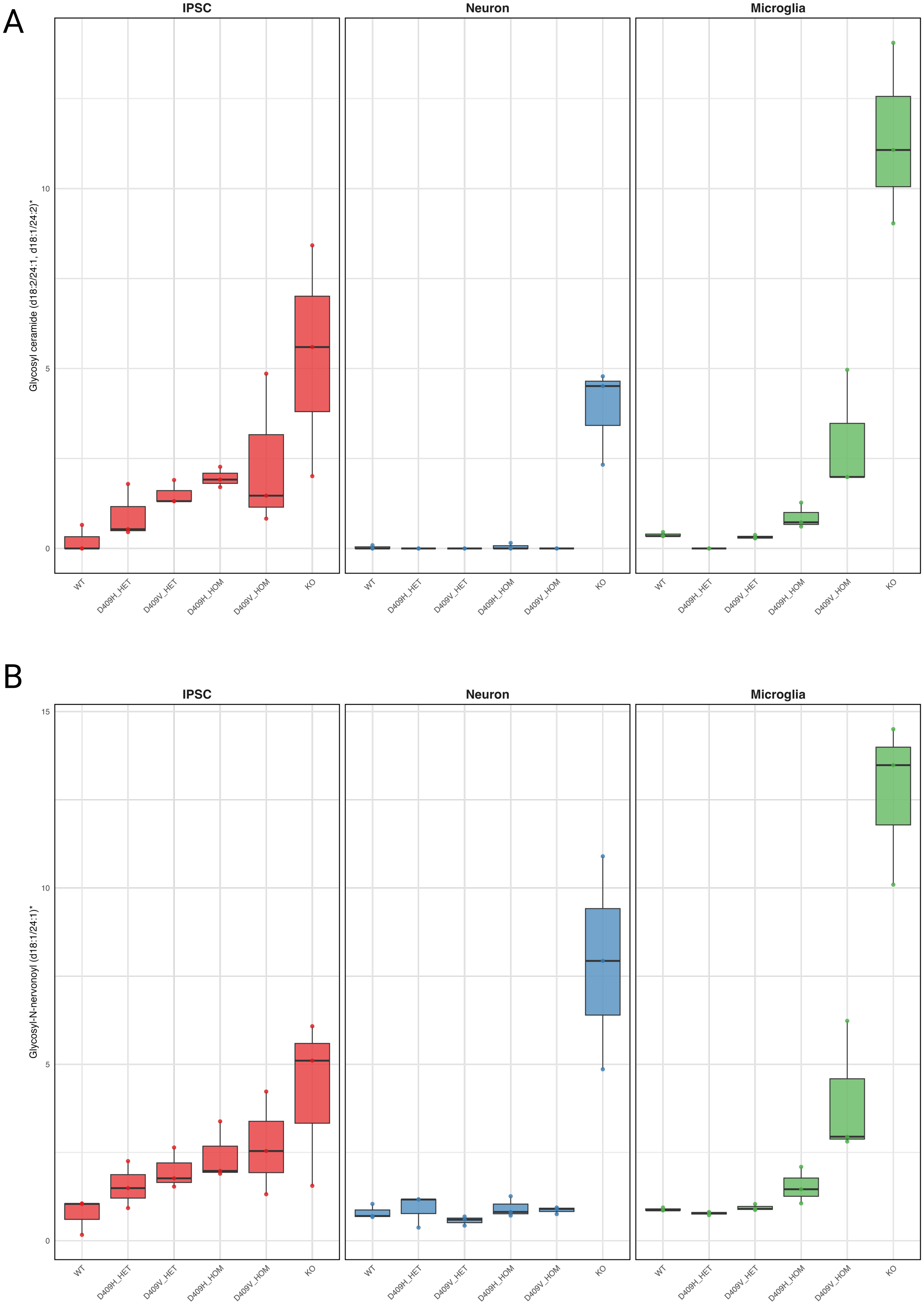
